# Mechanisms and physiological function of daily haemoglobin oxidation rhythms in red blood cells

**DOI:** 10.1101/2021.10.11.463714

**Authors:** Andrew D. Beale, Edward A. Hayter, Priya Crosby, Utham K. Valekunja, Rachel S. Edgar, Johanna E. Chesham, Elizabeth S. Maywood, Fatima H. Labeed, Akhilesh B. Reddy, Kenneth P. Wright, Kathryn S. Lilley, David A. Bechtold, Michael H. Hastings, John S. O’Neill

## Abstract

Cellular circadian rhythms confer temporal organisation upon physiology that is fundamental to human health. Rhythms are present in red blood cells (RBCs), the most abundant cell type in the body, but their physiological function is poorly understood. Here, we present a novel biochemical assay for haemoglobin (Hb) oxidation status which relies on a redox-sensitive covalent haem-Hb linkage that forms during SDS-mediated cell lysis. Formation of this linkage is lowest when ferrous Hb is oxidised, in the form of ferric metHb. Daily haemoglobin oxidation rhythms are observed in RBCs cultured *in vitro* or taken from freely behaving mice or humans exhibit and are unaffected by mutations that affect circadian rhythms in nucleated cells. These rhythms correlate with daily rhythms in core body temperature, with temperature lowest when metHb levels are highest. Raising metHb levels with dietary sodium nitrite can further decrease daytime core body temperature in mice via NO signaling. These results extend our molecular understanding of RBC circadian rhythms and suggest they contribute to the regulation of body temperature.

## Introduction

Daily rhythms in behaviour and physiology are observed in all kingdoms of life and are of fundamental importance for understanding human health and disease (Patke *et al*, 2020). In mammals, the daily organisation of cellular homeostasis occurs through the interaction between cell-intrinsic timing mechanisms with daily systemic cues; giving rise to rhythms in protein activity, electrical excitability, and cell motility, for example (Stangherlin *et al*, 2021). Cell-autonomous circadian timing can be affected by mutations in a number of proteins including kinases (e.g. CK1, CK2 and GSK3), transcription factors (e.g. CLOCK and BMAL1), and repressors such as PER and CRY (Ko & Takahashi, 2006). In most cell types, whilst the identity of ‘clock-controlled genes’ varies with tissue context, the rhythmic regulation of clock-controlled transcription is proposed to be the central cell-intrinsic mechanism by which cellular biology and function manifests as a daily rhythm (Ruben *et al*, 2018; Zhang *et al*, 2014).

Circadian oscillations in cellular biology exist in the naturally anucleate red blood cell (RBC), however, and so cannot be attributable to rhythms in nascent transcription. Circadian regulation of metabolism, redox balance, proteasomal degradation, and membrane electrophysiology have all been observed in isolated RBCs (Ch *et al*, 2021; Henslee *et al*, 2017; O’Neill & Reddy, 2011; Homma *et al*, 2015; Cho *et al*, 2014). The period of these circadian rhythms is sensitive to inhibition of proteasomal degradation and the activity of casein kinase 1, as in nucleated cells (Beale *et al*, 2019; Cho *et al*, 2014). In the absence of transcription or translation, RBC circadian rhythms are hypothesised to reflect a post-translational oscillator (PTO) that involves CK1, a ubiquitous component of circadian rhythms across the eukaryotic lineage (O’Neill *et al*, 2020; Causton *et al*, 2015). RBCs may therefore serve as a tractable model for interrogation of the putative PTO mechanism and post-translational rhythmic regulation of cellular processes more generally (Wong & O’Neill, 2018).

Observations of circadian rhythms in RBCs raise two important questions. First, since anucleate RBCs derive from nucleated precursors during erythropoiesis, do TTFL-regulated rhythms in erythroid progenitors influence timekeeping in isolated RBCs once they are terminally differentiated and anucleate? Second, in what way might circadian rhythms in RBCs impact upon critical physiological functions, such as gas transportation? Here, we address these questions in three ways: (1) characterising a novel rhythmic process in RBCs to aid investigation of RBC circadian mechanisms; (2) using this tool to describe oscillations in RBCs derived from mice harbouring well-characterised circadian mutations and free-living human subjects; and (3) by pharmacological manipulation of rhythms to assess functional relevance. Taken together, we demonstrate a novel biomarker for circadian phase *in vivo* that may be relevant for understanding daily variation in oxygen delivery, peripheral blood flow, and body temperature.

## Results

### An additional circadian marker in RBCs

Circadian rhythms of NAD(P)H concentration, PRX-SO2/3 abundance and membrane physiology are observed in isolated mammalian red blood cells (Cho *et al*, 2014; Henslee *et al*, 2017; O’Neill & Reddy, 2011). While investigating PRX-SO2/3 rhythms in isolated human red blood cells, we noticed faint chemiluminescent bands at ∼16 kDa and ∼32 kDa which exhibited more robust circadian rhythmicity than PRX-SO2/3 (O’Neill & Reddy, 2011). This uncharacterised, rhythmic chemiluminescence was readily observed upon incubating membranes with enhanced chemiluminescence (ECL) reagent immediately after transfer from polyacrylamide gel, without any antibody incubations or exogenous source of peroxidase activity (Figure 1A, Expanded View figure 1A). Recognising the utility of an additional marker for RBC circadian rhythms, we sought to characterise the source of this chemiluminescence.

**Figure 1.**
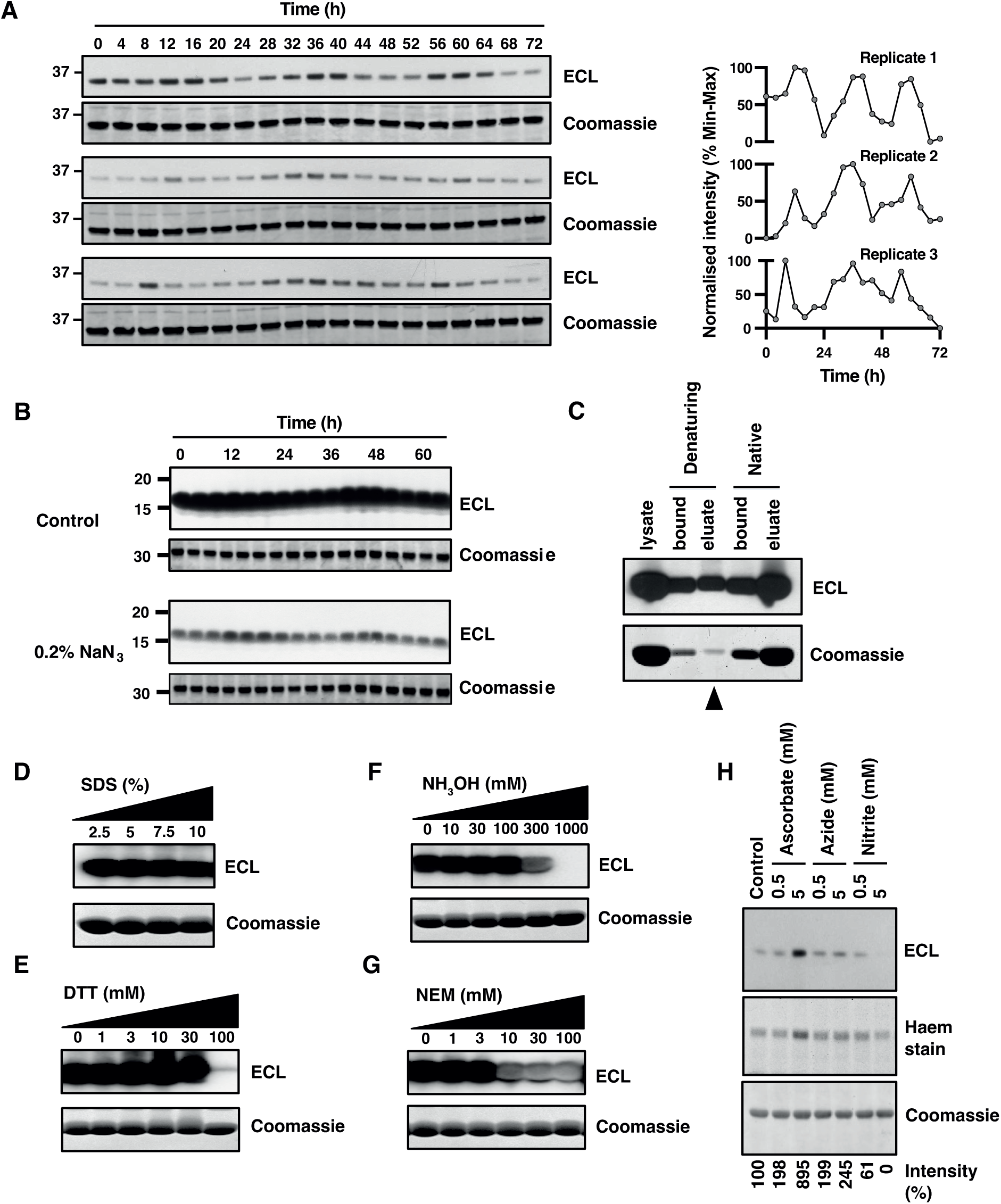
Biochemical characterization of a protein-bound peroxidase activity in isolated RBCs following detergent lysis and SDS-PAGE. **A** Circadian rhythm of peroxidase activity in 3 independently isolated sets of human red blood cells sampled every 4 hours under constant conditions, quantifications shown on right hand side (*n*=3). After lysis, SDS-PAGE and transfer to nitrocellulose, membranes were immediately incubated with ECL reagent to reveal peroxidase activity bound to the membrane. For practical reasons, the upper ∼32 kDa band (corresponding to the dimer, Hb2*) was used for quantification since the lower band quickly saturated the x-ray film used to detect chemiluminescence. Coomassie stained gels were used as loading controls; the ∼16 kDa band (corresponding to the monomer, Hb) from the coomassie stained gel is shown. **B** Representative blots performed in parallel following SDS-PAGE of human erythrocyte time course samples and transfer to nitrocellulose membranes. Before incubation with ECL reagent, membranes were incubated for 30 minutes in PBS ± 0.2% sodium azide. Coomassie stained gels were used as loading controls; the Hb band from the coomassie stained gel is shown. **C** Ni-NTA affinity purification of human RBC lysates under denaturing and native conditions. The indicated band (arrowhead) was excised for mass spectrometry. **D-F** RBC samples were lysed with a buffer containing increasing concentrations of (D) SDS; (E) DTT; or (F) hydroxylamine (NH3OH, pH 7) and incubated for 30 minutes at room temperature. Samples were incubated with ECL reagent following SDS-PAGE and transfer to nitrocellulose. **G** Intact RBCs were incubated with increasing concentrations of N-ethylmaleimide (NEM) for 30 minutes at room temperature before lysis. Samples were incubated with ECL reagent following SDS- PAGE and transfer to nitrocellulose. **H** RBCs were pre-incubated for 30 minutes in Krebs buffer at room temperature containing sodium ascorbate, sodium azide, sodium nitrite or sodium chloride (5 mM) as a control and subject to SDS- PAGE and nitrocellulose transfer. Representative ECL-only blot and in-gel haem staining is shown for the upper, Hb2 band. ECL staining intensity relative to control is shown. In all panels, the Hb band from coomassie stains of the protein remaining in gels subsequent to transfer onto nitrocellulose membranes is shown as loading control.

The presence of chemiluminescence in the absence of antibody suggested that some RBC protein species possess an intrinsic peroxidase activity that is tolerant to the denaturing buffer (containing 1% dodecyl sulphate pH 8.5) used for cell lysis. The ECL peroxidase reaction employed in modern immunoblotting is usually catalysed by horseradish peroxidase but can, in fact, be catalysed by any haem group (Das & Hecht, 2007; Keilin & Hartree, 1950). At high concentrations, sodium azide (NaN3) inactivates peroxidase activity of haem groups through azidyl radical addition to the prosthetic group, and is a common means of inactivating peroxidase activity in immunoblotting (Montellano *et al*, 1988). To confirm that this SDS-resistant species indeed catalyses the ECL reaction through an intrinsic peroxidase activity, we incubated freshly transferred membranes with 30 mM (0.2%) NaN3 for 30 minutes prior to assaying peroxidase activity *via* ECL (Figure 1B).

Pre-treatment with azide elicited a marked reduction in chemiluminescence, indicating that a haem-mediated (non-HRP) peroxidase activity was indeed adsorbed to the nitrocellulose membrane following transfer after SDS-PAGE. Haemoglobin (Hb, MW 16 kDa) is the most common haem protein in RBCs, comprising 97% of all RBC protein (Pesciotta *et al*, 2015; Roux-Dalvai *et al*, 2008) with a small minority of Hb monomers being observed to run as a cross-linked dimer (Hb2, ∼32 kDa) by SDS-PAGE (Fuhrmann *et al*, 1988). We therefore considered it plausible that the rhythmic non-specific peroxidase activity might be attributable to a covalent haem-Hb moiety: a linkage that would be intrinsically resistant to denaturation *via* SDS and subsequent polyacrylamide gel electrophoresis. In RBCs under physiological conditions, however, Fe-haem is well characterised as being bound to Hb protein through non-covalent co-ordination by proximal and distal histidine residues (Perutz *et al*, 1960). Therefore, it should not be resistant to denaturation by SDS. Importantly, however, covalent linkage of haem to haem-binding proteins, including globins, has been reported to occur under several non-physiological conditions (Catalano *et al*, 1989; Deterding *et al*, 2004; Enggist *et al*, 2003; Reeder *et al*, 2008; Reeder, 2010).

To assess whether some fraction of Fe-haem is covalently bound to Hb protein, we employed Ni-NTA affinity chromatography under native and denaturing conditions. Under native conditions, haem proteins are readily bound by Ni-NTA (Pesciotta *et al*, 2015). Under denaturing conditions, we reasoned that only proteins that are covalently linked to haem should show high affinity for the Ni-NTA matrix. Commensurately, we found that the non-specific peroxidase bound to, and eluted from, Ni-NTA under both native and denaturing conditions (Figure 1C). Mass spectrometry of the eluted 16 kDa protein revealed the major species to be haemoglobin beta and alpha ( ≥ 95% coverage) (Expanded View figure 2A). Moreover, mass spectrometry of full length HbA, purified under denaturing conditions, revealed additional peaks at +614 Da compared with the HbA polypeptide, which corresponds to the mass of deprotonated haem b (Expanded View figure 2B). Therefore, the SDS-resistant peroxidase activity (Figure 1D) was assigned to haemoglobin.

We considered that the unsaturated bonds in the haem porphyrin ring should be an attractive target for Click chemistry through base-catalysed Michael addition from a cysteinyl thiol under the denaturing and basic RBC lysis conditions we employ (Nair *et al*, 2014). This reaction would be expected to produce a thioester-linked adduct of Fe-haem to Hb protein, though likely with a poor yield since this reaction would be competing with mixed disulphide bond formation under these lysis conditions. To test this, we included high concentrations of strong reductants (DTT and NH3OH) in the lysis buffer, which reduce thioester (but not thioether) bonds under aqueous conditions (Pedone *et al*, 2009). We found that the residual peroxidase activity of Hb on nitrocellulose membranes was completely abolished under these conditions (Figure 1E, F), supporting a thioester linkage. To validate this, we pre-incubated RBCs with N-ethylmaleimide (NEM) to alkylate Hb cysteine residues prior to lysis - effectively blocking *de novo* thioester bond formation without affecting any thioester bonds that might exist prior to lysis. NEM pre-treatment abolished subsequent peroxidase activity (Figure 1G), indicating that thioester bond formation is facilitated by protein denaturation during cell lysis.

Overall, our observations accord with a model whereby the rhythmic peroxidase activity we detect arises from a small proportion of Hb becoming thioester-linked to haem during cell lysis, but does not explain why the formation of this bond might exhibit a circadian rhythm. The total amount of Hb detected by coomassie was invariant throughout the RBC circadian cycle (Figure 1A, Expanded View figures 1, 3); whereas the peroxidase activity associated with Hb (and Hb2) showed a clear circadian rhythm (Figure 1A, Expanded View figures 1, 3). Moreover, the proportion of Hb protein that was covalently linked to haem-Hb was very low compared with the total amount of RBC Hb (Figure 1C). Previous work has indicated that RBCs exhibit a circadian rhythm in the oxidation of certain protein species including the oxidation and over-oxidation of peroxiredoxin proteins, the latter being degraded by the proteasome (Cho *et al*, 2014), as well as rhythms in Hb-quaternary structure and the cellular reducing equivalents (NAD(P)H), that are normally used by methaemoglobin (metHb) reductase to reduce metHb back to the ground state (O’Neill & Reddy, 2011). These are suggestive of a circadian rhythm in the oxidation of oxyhaemoglobin to form ferric (inactive, deoxy, Fe(III)) metHb, and subsequent H2O2 formation, which possibly results as a consequence of daily variation in Hb dimer-tetramer equilibrium (O’Neill & Reddy, 2011).

In light of these observations, and the strong evidence for dynamic regulation of Hb redox state in RBCs under physiological conditions (Umbreit, 2007), we asked whether the redox state of haem-Hb at the point of cell lysis might affect the level of thioester bond formation. Deoxygenated ferrous (Fe(II)) Hb is readily susceptible to oxidation to ferric (Fe(III)) metHb by nitrite at low millimolar concentrations that will not oxidise cysteine (Cortese-Krott *et al*, 2015). Conversely, millimolar ascorbate reduces metHb back to the ferrous state (Eder *et al*, 1949; Gibson, 1943), whereas similar concentrations of azide, a less favourable electron donor or acceptor, should have less effect on Hb redox state under these conditions. Acute treatments of intact RBCs prior to lysis with sodium nitrite and sodium ascorbate dose-dependently decreased and increased, respectively, the peroxidase activity detected at Hb molecular weight on nitrocellulose membranes (Figure 1H), compared with a much more modest effect of sodium azide. An in-gel colourimetric haem stain yielded similar results to the ECL assay (Figure 1H).

Taken together, our observations support a model whereby a small proportion of ferrous haem spontaneously crosslinks with Hb via a thioester bond upon cell lysis which is stable during subsequent SDS-PAGE and transfer to nitrocellulose. This proportion is influenced by the redox state of haem-Hb in the RBCs prior to lysis and hence lysis-induced crosslinking reveals the underlying redox status of Hb. The Hb-based marker demonstrated here, allowing detection of RBC circadian rhythms by ECL reagent alone, represents a novel report for RBC circadian rhythms, a second facet of the redox rhythm in RBCs. We coin this technique “Bloody Blotting”, distinguishable from immunoblotting by the lack of any antibodies, blocking step or exogenous source of peroxidase activity.

### Clock mutations do not affect RBC circadian rhythms

Oscillations in cellular processes in RBCs cannot be attributable to transcriptional repression by PER and CRY proteins since these proteins are not present in RBCs, nor do RBCs possess the capacity for transcriptional feedback (Bryk & Wiśniewski, 2017; O’Neill & Reddy, 2011). Consistent with this, circadian rhythms in mature isolated RBCs are also insensitive to inhibition of nascent transcription and translation (O’Neill & Reddy, 2011). However, the RBC develops from nucleated precursors, the normoblasts, during erythropoiesis and we therefore considered whether the developmental expression of circadian cycles in erythroid precursors might affect circadian phenotype of mature RBCs. To address this possibility, we isolated RBCs from mice harbouring well-characterised circadian mutations of the post-translational regulators of PER and CRY, CK1;*^Tau/Tau^*(Lowrey *et al*, 2000; Meng *et al*, 2008) and FBXL3*^Afterhours/Afterhours^*(Godinho *et al*, 2007) respectively, and examined circadian rhythms under constant conditions using PRX-SO2/3 abundance and Bloody Blotting as rhythmic reporters.

Mice homozygous for these mutations exhibit behavioural circadian periods that are shorter (*tau* mutant) and longer (*afterhours*, *afh* mutant) than wildtype (Godinho *et al*, 2007; Lowrey *et al*, 2000; Meng *et al*, 2008). Since circadian timekeeping occurs cell-autonomously, the short and long period phenotypes observed at the whole organism level are readily recapitulated in fibroblasts isolated from homozygous mutant *Cry1:luciferase* mice (Figure 2A). Fibroblasts harbouring *tau* or *afh* mutations exhibit circadian periods 2.4 h shorter and 6.7 h longer, respectively, than wildtype (Figure 2A). In RBCs isolated from the same mice and cultured *ex vivo*, significant oscillations were observed in PRX-SO2/3 abundance (Figure 2B, D) and Hb peroxidase activity (Hb2*, Figure 2C, E, Expanded View figure 3A, B, C). However, unlike fibroblasts, no significant difference in circadian period was seen between the three mouse genotypes (Figure 2F). Therefore, the altered timing observed in mutants of nucleated cells is not inherited by RBCs and the timekeeping role of CK1 in RBCs (Beale *et al*, 2019) is within a PTO that does not involve PER, CRY, CK1ε or FBXL3 (Bryk & Wiśniewski, 2017; O’Neill & Reddy, 2011). Furthermore, the similar circadian period between these reporters of redox state, irrespective of genotype origin, indicates they are two outputs of the same underlying oscillation, but with a higher relative amplitude and earlier phase in the Hb2* bands compared with PRX-SO2/3.

**Figure 2.**
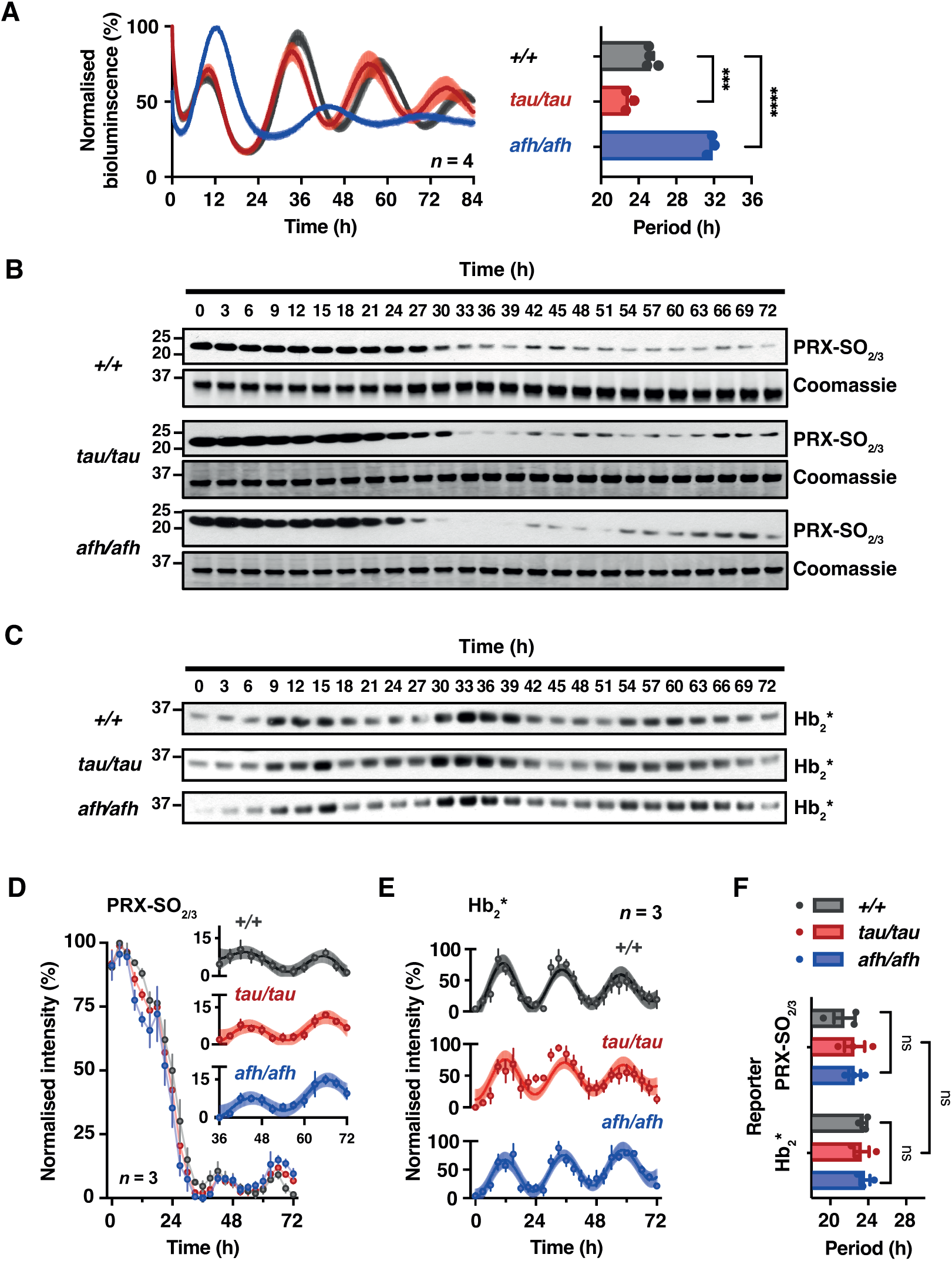
The Clock in Mouse Erythrocytes is Unaffected by Mutations that Affect Circadian Period in Nucleated Cells. **A** Primary fibroblasts isolated from CK1ε*^Tau/Tau^* or Fbxl3a*^Afh/Afh^* mutant mice, also transgenic for the *Cry1:luc* reporter, exhibit bioluminescence rhythms that are shorter and longer than wild type control, respectively. Normalised mean±SEM are shown as line and shading respectively (*n*=4). Grouped quantification of circadian period from A. p<0.0001 by 1-way ANOVA; Sidak’s multiple comparisons test displayed on graph. **B** RBCs from CK1ε*^Tau/Tau^*, Fbxl3a*^Afh/Afh^* and wild type mice were incubated at constant 37°C from the point of isolation. Single aliquots were lysed every three hours. Representative immunoblot showing PRX monomer over-oxidation over 72 hours under constant conditions. Coomassie stained gels were used as loading controls; the Hb band from the coomassie stained gel is shown. **C** Representative membranes showing peroxidase activity detected without antibody following transfer to nitrocellulose membrane, at the same molecular weight as haemoglobin dimer (∼32 kDa), indicated as Hb2*. Loading control is shown in (B) by coomassie stain. **D** Grouped quantification from (B) of PRX-SO2/3 oscillation (*n*=3 per genotype). 2-way ANOVA, p<0.0001 for time effect, but not significant for genotype or time/genotype interaction (p>0.05); inset shows the final 36 hours with expanded y-axis. Damped cosine wave using least squares non-linear regression fitted to final 36 hours shown as line, with SEM error shown as shading. **E** Grouped quantification from (D) of Hb2* oscillation (*n*=3 per genotype). 2-way ANOVA, p<0.0001 for time effect, but not significant for genotype or time/genotype interaction (p>0.05). **F** Circadian periods of PRX overoxidation, PRX-SO2/3, and peroxidase activity, Hb2*, derived from damped cosine fits from (D) and (E) respectively. No significant difference by 2-way ANOVA. Data information: In (A-F), data are presented as mean ± SEM, *p≤0.05, **p≤0.01, ***p≤0.001 , ****p≤0.0001.

### Daily variation in redox status in humans *in vivo*

*In vivo*, haem-Hb occurs in human RBCs in two oxidation states: as ferrous Hb(II) and as ferric metHb(III). The transport of oxygen requires oxygen reversibly bound to ferrous Hb(II). Oxygenated tetrameric Hb(Fe(II))O2 is a very stable complex but does slowly auto-oxidize at a rate of about 3-4 %/day (Eder *et al*, 1949; Johnson *et al*, 2005) a rate that is accelerated at lower partial pressures of O2 (pO2) when the haemoglobin is partially oxygenated. In a healthy individual, metHb reductase ensures that < 1 % of RBC Hb exists in the Fe(III) state, although disease, dietary nitrites and inherited conditions such as methaemoglobinemia can elevate this (Cawein *et al*, 1964). Given the daily variation of Hb redox status suggested here and in previous studies in mouse and human RBCs *in vitro* (Cho *et al*, 2014; O’Neill & Reddy, 2011), we used “Bloody Blotting” to determine whether daily variation in Hb(II):metHb(III) ratio occurs *in vivo*. We therefore collected and flash froze blood samples every 2 hours from 4 healthy volunteers, beginning at habitual wake time, over a complete circadian cycle under controlled laboratory conditions (Figure 3A). We observed a significant variation in Hb- linked peroxidase activity that peaked shortly after waking and reached its nadir 12 hours later (Figure 3B and C, Expanded View figure 1B). This, together with our biochemical and *in vitro* data, suggests that the Hb(II):metHb(III) ratio varies with time of day, with the proportion of metHb in the blood highest (peroxidase activity lowest) towards the end of the habitual waking period.

**Figure 3.**
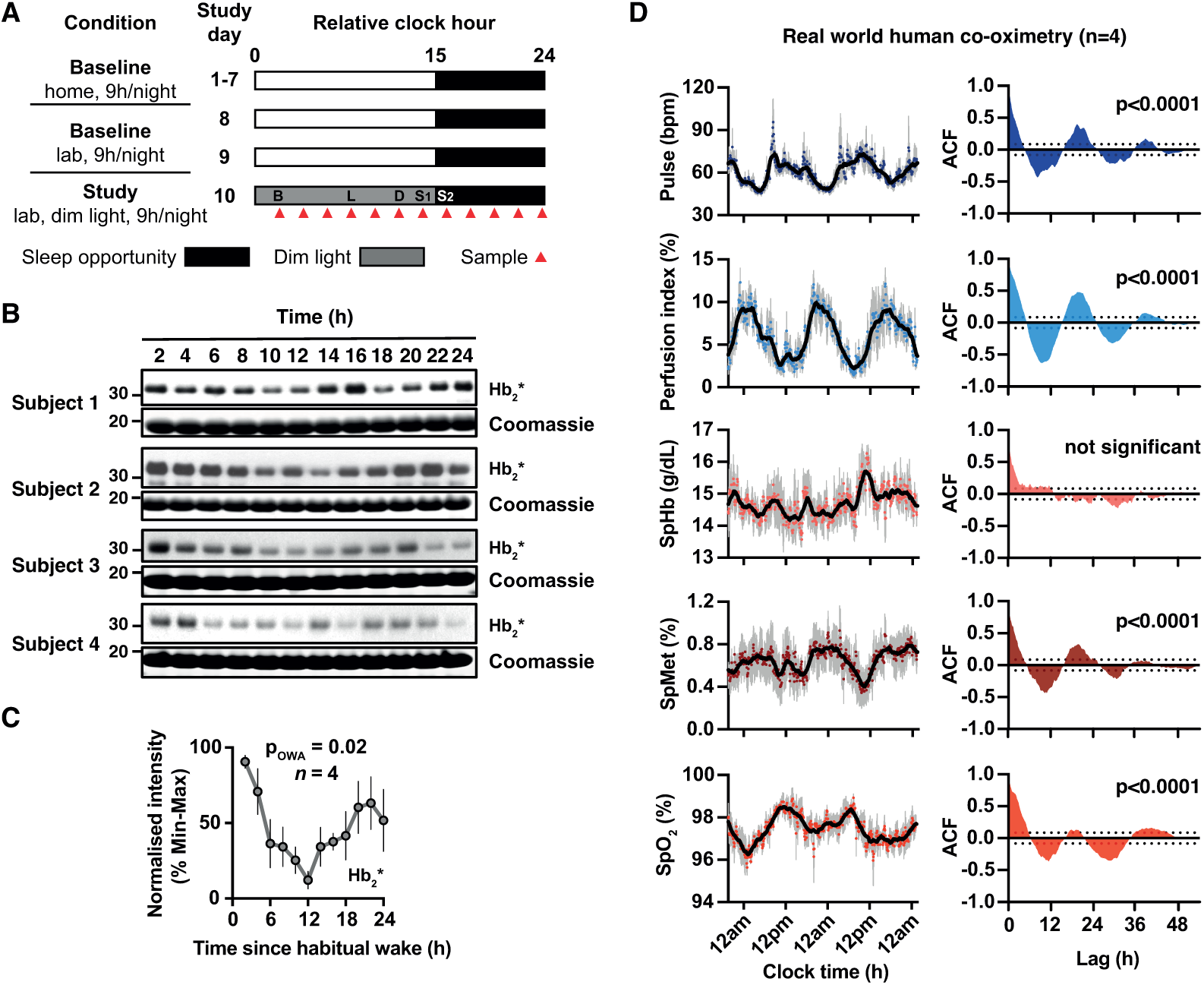
**Daily variation in redox status in humans *in vivo*** **A** Experimental protocol for four healthy participants, relative to waketime at 0. Participants maintained consistent 15h:9h days consisting of wake (white) and sleep opportunity (black) at home for at least one-week prior to entering the laboratory. Following two baseline laboratory days, participants were awakened at habitual waketime and studied under dim light conditions (< 10 lux in the angle of gaze during wakefulness and 0 lux during scheduled sleep), given an energy balanced diet (standardised breakfast, B, lunch, L, and dinner, D, meals given at 2h, 8h and 12h respectively, and two after dinner snacks, S1 and S2, at 14h and 16h, water available *ad libitum*). Blood samples were collected every two hours (red triangles). Daytime and light exposure during scheduled wakefulness is depicted as white; night-time and sleep opportunity is depicted as black; dim light during wakefulness is depicted as dark grey. **B** Whole blood samples from 4 volunteers were subject to SDS-PAGE and immunoblot. Representative Hb2* peroxidase activity (upper). Coomassie stained gels were used as loading controls; the Hb band from the coomassie stained gel is shown. **C** Quantification of 4 subjects. One way ANOVA, p<0.05. **D** Pulse co-oximetry data from four healthy male volunteers under normal daily life (*n*=4). (Left) Pulse rate, perfusion index, total haemoglobin concentration SpHb, methaemoglobin proportion SpMet, and oxygen saturation SpO2 are plotted against local time (coloured dots = grouped means; grey lines = error bars; solid black line = 1-hour moving average). (Right) Autocorrelation function (ACF) against time lag (h) of the grouped means. 95% confidence bands of the ACF are plotted as dotted lines on y-axis, p value represents extra sum-of-squares F test comparing a straight line fit (H0) with a damped cosine fit. Data information: In (C), data are presented as mean ± SEM; in (D), data are presented as mean ± SEM with individual points. *p≤0.05, **p≤0.01, ***p≤0.001 , ****p≤0.0001.

To test whether daily variation in metHb levels occurs in a real-world setting, we assessed metHb levels and other blood parameters in 4 free-living healthy human subjects using pulse co-oximetry. As expected, pulse rate (PR) exhibited a clear daily variation in these subjects (Figure 3D, left), with significant autocorrelation in the circadian range (Figure 3D, right). Perfusion index (PI, a measure of peripheral blood flow) (Goldman *et al*, 2000) also exhibited a clear daily variation, peaking around midnight, approximately antiphasic to the daily variation in pulse rate. Remarkably, in contrast to total Hb (SpHb) that displayed no significant 24h variation, the proportion of metHb (SpMet) in the blood exhibited a striking daily variation that rose during the evening and peaked during the night (Figure 3D). These subjects were in a real-world setting, and thus affected by environmental and social cues from a normal working day. However, the evening rise and night-time peak is consistent with the observed reduction in Hb2* activity at the end of the waking period in laboratory conditions (Figure 3B). The oxygen saturation of the blood (SpO2) varied in antiphase with the metHb rhythm, peaking during the active phase (Figure 3D), consistent with the previously established relationship between Hb oxygenation and metHb formation (Umbreit, 2007). This suggests one functional consequence of the metHb rhythm is a daily variation in the oxygen carrying capacity of blood.

### Effect of rhythms in metHb on vascular flow and body temperature

In addition to the possible effect on the oxygen-carrying capacity of the blood, a second consequence of daily cycles of Hb redox status is the regulation of vascular tone and peripheral blood flow through nitric oxide (NO) signalling (DeMartino *et al*, 2019; Grubina *et al*, 2007; Huang *et al*, 2005; Umbreit, 2007). RBCs store, carry and release NO equivalents (nitrite, nitrosyl adducts, and N2O3 respectively) that complement tissue-resident local NO synthesis to stimulate peripheral vasodilation (Basu *et al*, 2007; Grubina *et al*, 2007; Cosby *et al*, 2003). MetHb reacts with nitrite to yield an intermediate that rapidly reacts with NO to yield N2O3, which is proposed as the essential vector for release of NO from RBCs (Basu *et al*, 2007). We note that increased peripheral blood flow is the primary mechanism allowing heat release from the core and, in humans, is essential for lowering core body temperature at night that habitually accompanies the transition from wake to sleep (Aschoff & Heise, 1974; Smolander *et al*, 1993; Murphy & Campbell, 1997; Rzechorzek *et al*, 2022). Thus, increased metHb during the rest phase (at night in humans, in the day in mice) should contribute to increased peripheral blood flow (Figure 3D), and consequently the reduction in core body temperature (Figure 4A).

**Figure 4.**
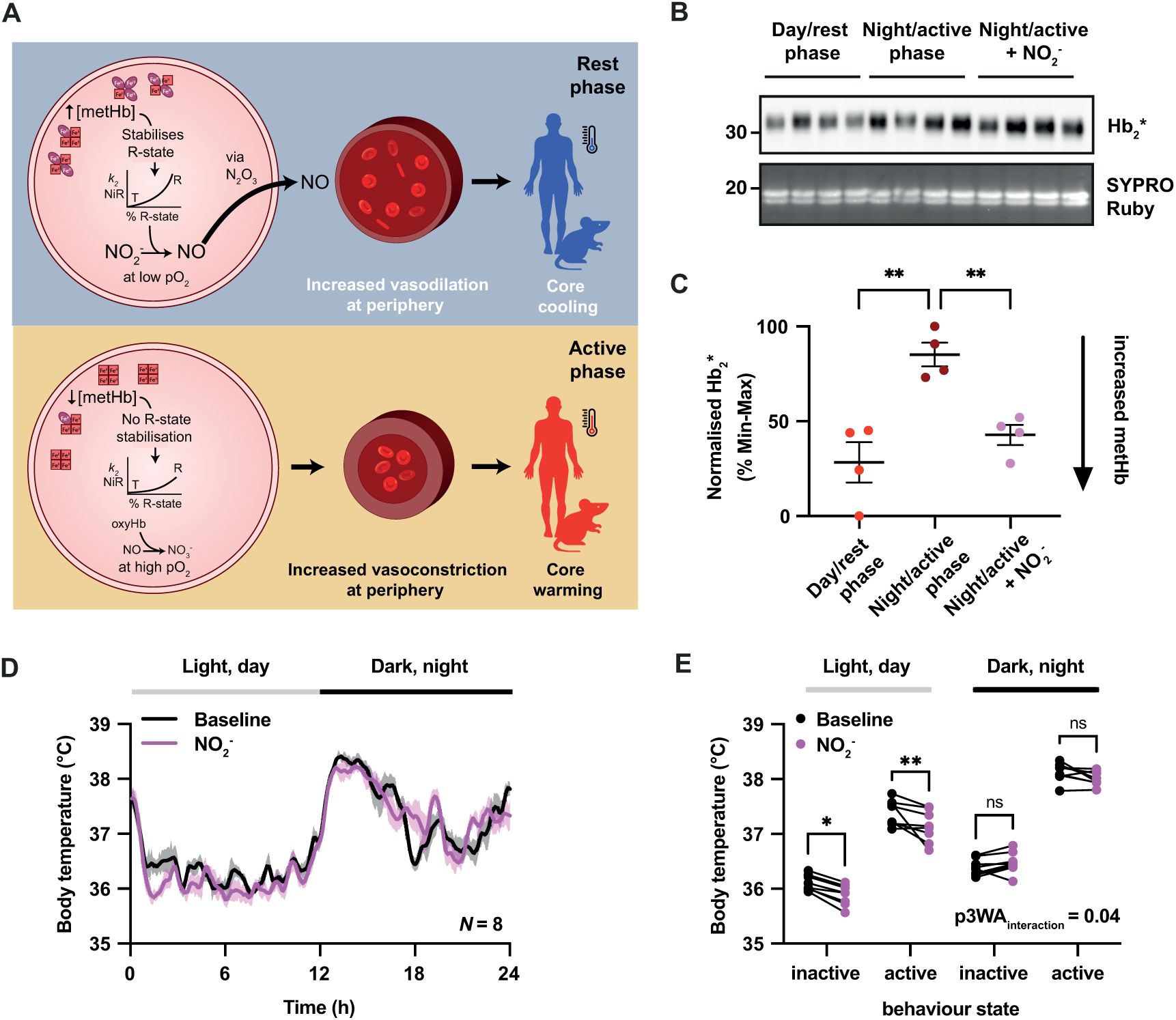
Perturbation of metHb levels *in vivo* affects body temperature via NO signalling. **A** A hypothesis for the relationship between metHb levels and body temperature. In the rest phase (night for humans; day for mice), increasing stabilisation of the R state of haemoglobin as levels of metHb rise (Bunn & Forget, 1986) leads to an increase in affinity for O2 (Monod *et al*, 1965; Perutz, 1970) and an increased nitrite reductase activity of deoxyHb at low pO2 (Grubina *et al*, 2007; Huang *et al*, 2005). The subsequent export of increased concentrations of NO, via an N2O3 intermediate (Basu *et al*, 2007), leads to increased vasodilation (Cosby *et al*, 2003; Crawford *et al*, 2006) and thus increased heat loss. In the active phase, the concentration of metHb is lower, reducing deoxyHb nitrite reductase activity and thus RBC-produced NO. When fully oxygenated, oxyHb inactivates NO, or NO2^-^, to nitrate, NO3^-^ (Herold & Shivashankar, 2003), reducing vasodilation. Taken together, the daily variation in metHb and its effect on the production and export of NO, and subsequent vasodilation, is predicted to lead to a daily variation in body temperature due to daily variation in vasodilation- mediated cooling (Krauchi & Wirz-Justice, 1994). **B** Whole blood samples were taken from mice during the early rest phase (day, ZT3), early active phase (night, ZT15), or during the early active phase from mice which received supplemental dietary nitrite (50mg/l) for 10 days. Whole blood was subject to SDS-PAGE and immunoblot. Hb2* peroxidase activity (upper) and Hb band from SYPRO Ruby blot staining serves as a loading control (lower). **C** Quantification of normalised Hb2* activity from 4 mice. Mean±SEM, p=0.0007 by 1-way ANOVA; Sidak’s multiple comparisons test displayed on graph. **D** Average daily body temperature profile from mice ± dietary nitrite over 2 days in light (day, light grey bar) and dark (night, black bar) cycles. *N*=8 mice per treatment condition. **E** For each mouse in each treatment condition (*n*=8 mice per condition), average body temperature was calculated during inactive (activity < median activity across 2-day recording) and active (activity > median activity across 2-day recording) periods in light and dark phases. Thus, for each mouse, four separate average body temperatures were generated, belonging to inactive and active periods during the light phase, and inactive and active periods during the dark phase. Repeated measures 3-way ANOVA ptreatment < 0.01, pactive_vs_inactive < 0.0001, pday_vs_night < 0.0001, pinteraction: inactive vs active x baseline vs nitrite < 0.05; Sidak’s multiple comparisons test for interaction: inactive vs active x baseline vs nitrite displayed. Data information: In (C-D), data are presented as mean ± SEM; in (E), data are presented as individual, paired, points. *p≤0.05, **p≤0.01, ***p≤0.001 , ****p≤0.0001.

Like humans, mice exhibit significant daily rhythms in body temperature that arise from a 24h variation in the mismatch between heat generation and heat loss (Refinetti & Menaker, 1992a). Daily rhythms of heat generation over the daily cycle have been intensively studied, with nocturnal increases in locomotor activity and brown fat thermogenesis being dominant factors in rodents (Refinetti & Menaker, 1992a; Cannon & Nedergaard, 2004). The contribution of increased peripheral vasodilation, and therefore heat loss, during the rest phase to the daily rhythm in body temperature has received less attention.

To test the prediction that increased metHb levels during the rest phase contribute to heat loss (Figure 4A), we used sodium nitrite, a direct modifier of Hb oxidation status *in vitro* (Figure 1H), a common cause of methaemoglobinemia in humans (Dela Cruz *et al*, 2018), and a known vasodilator *in vitro* (Ignarro *et al*, 1981) and *in vivo* (Cosby *et al*, 2003). We first validated the feasibility of direct metHb perturbation *in vivo*, by providing 50 mg/L sodium nitrite to mice in drinking water over 7 days, assessed by Bloody Blotting (Figure 4B, Expanded View figure S4). Like human blood taken *in vivo* (Figure 3C), Hb2* activity was significantly higher in mice sampled during the early active phase (night) than early rest phase (day) (Figure 4B). When given sodium nitrite, blood Hb2* activity collected in the active phase was significantly reduced relative to untreated mice (Figure 4B) in accordance with *in vitro* data (Figure 1H) and the prior literature. This confirms that nitrite can indeed be used to manipulate metHb *in vivo*.

In mice we expect core body temperature to be lower during the daytime, when animals consolidate their sleep, than at night, and predicted that this difference would be accentuated by nitrite. We tested this in mice implanted with telemetric activity and temperature sensors: boosting metHb levels by providing nitrite in drinking water, while correcting for daily differences in locomotor thermogenesis by considering activity state as a separate variable. Consistent with prediction, we found a significant daytime-specific decrease in core body temperature, both at rest and when mice were physically active, when mice were treated with nitrite (Figure 4 D,E).

Taken together, our results establish a novel reporter for circadian rhythms in RBCs *in vitro* and provide further insight into the determinants and physiological outputs of the circadian clock in the most abundant cell of the human body (Alberts *et al*, 2017; Sender *et al*, 2016). This novel reporter is dependent on the redox status of Hb, which alters the proportion of haem that is covalently linked to Hb on cell lysis. The Hb oxidation status is under circadian control *in vitro* and *in vivo* but is independent of the TTFL-based timing mechanism of RBC precursors. Finally, we show that the redox status of Hb may have a direct impact on core body temperature via NO-signalling and vasodilation, and is the first indication of a functional consequence for RBC circadian rhythms.

## Discussion

RBCs have been an interesting model for circadian rhythms in the absence of transcription for a number of years. However, the functional relevance of these rhythms to RBCs has remained elusive. Here we have extended understanding of RBC circadian function, and uncovered a novel RBC clock marker based upon a redox-sensitive covalent haem-Hb linkage that forms during cell lysis/protein denaturation. This “Bloody Blotting” is fast and inexpensive; however, its utility as an RBC phase marker *in vivo* is likely limited to research contexts. In contrast, pulse co-oximetry, a wearable technology, accurately reports physiological diurnal variation in relevant, haemodynamic parameters simultaneously over many days. We show that RBC rhythms are independent of their developmental pathway and have functional significance in their O2-carrying and NO-generating capacity.

### Multiple reports point to a single oscillator in RBCs

Circadian rhythms have previously been described in isolated human and mouse RBCs (Beale *et al*, 2019; Cho *et al*, 2014; Henslee *et al*, 2017; O’Neill & Reddy, 2011; Ch *et al*, 2021) under comparable free run conditions to those employed here. However, those experiments employed *ex vivo* entrainment by applied 12h:12h temperature cycles to mimic body temperature rhythms, whereas here, we culture RBCs in constant conditions directly after isolation from the mouse. Despite the differences in entrainment (*ex vivo* vs *in vivo*), PRX-SO2/3 immunoreactivity in isolated RBCs consistently peaked at a point equivalent to the middle to end of the active phase *in vivo* (at the transition from hot to cold). MetHb levels peaked (Hb2* activity lowest) later, at the beginning of the rest phase in mouse RBCs *ex vivo* and humans *in vivo*. PRX-SO2/3 and Hb redox state oscillated with the same circadian period, as has been shown for rhythms in PRX-SO2/3 abundance, membrane physiology and central carbon metabolites (Ch *et al*, 2021; Henslee *et al*, 2017). This suggests that there is a single underlying molecular oscillator in RBCs, whose activity is dependent on ion transport, the 20S proteasome and CK1 (Beale *et al*, 2019; Cho *et al*, 2014; Henslee *et al*, 2017) and whose rhythmic outputs include Hb redox state, PRX-SO2/3 degradation, metabolic flux and K^+^ transport. As with other cell types, RBC rhythms *in vivo* are presumably synchronised by systemic cues (Dibner *et al*, 2010) which continue under constant routine conditions with respect to posture, food intake and sleep (Skene *et al*, 2018). Whilst we cannot rule out that systemic cues, or indeed body temperature or sleep-wake rhythms, are entirely responsible for metHb rhythms observed in humans *in vivo*, taken together with *ex vivo* data, our observations are consistent with the interpretation that RBC rhythms are under cell-autonomous circadian control and synchronised by systemic timing cues.

### Physiological Relevance

By mechanisms that remain to be firmly established, in isolated RBCs there is a circadian rhythm in the rate of Hb auto-oxidation and metHb formation (O’Neill & Reddy, 2011). Nevertheless, daily regulation of Hb redox status has functional significance in at least two ways. Since metHb cannot bind O2, metHb rhythms may affect the oxygen-carrying capacity of the blood. We found a daily rhythm of modest relative amplitude in SpO2 in antiphase with metHb (Figure 3D). Though this is correlational, the phase relationship between the rhythm in metHb and SpO2 is consistent with the influence of metHb on Hb’s allosteric Hill co-efficient. MetHb stabilises the R state of Hb (Gladwin & Kim-Shapiro, 2008) meaning that when metHb is higher, transition to the Hb T-state will occur at relatively lower pO2 and thereby facilitate increased oxygen supply to peripheral blood vessels/tissues during the rest phase. However, the low amplitude is likely to limit its the physiological significance for oxygen-carrying capacity.

A more important potential consequence of daily cycles of Hb redox status is the regulation of vascular tone and peripheral blood flow through nitric oxide (NO) signalling (DeMartino *et al*, 2019; Grubina *et al*, 2007; Huang *et al*, 2005; Umbreit, 2007). RBCs store, carry and release NO equivalents (nitrite, nitrosyl adducts, and N2O3 respectively) through RBC-dependent NO generation when pO2 falls (Basu *et al*, 2007; Grubina *et al*, 2007; Cosby *et al*, 2003), which complements local NO synthesis by tissue-resident nitric oxide synthases. MetHb reacts with nitrite to yield an intermediate that rapidly reacts with NO to yield N2O3, which is proposed as the essential vector for release of NO from RBCs to stimulate hypoxic vasodilation in all peripheral blood vessels (Basu *et al*, 2007). At high pO2, oxygenated Hb(II) inactivates local NO, resulting in vasoconstriction (Doyle & Hoekstra, 1981; Umbreit, 2007). Thus, based on current understanding of human physiology, increased metHb during the rest phase (at night) should contribute to increased peripheral blood flow. This can be clearly discerned from the high amplitude diurnal variation in perfusion index we observed (Figure 3D).

A second critical aspect of vasodilation lies in its contribution to regulating core body temperature. Heat generated during physical exercise and by mitochondrial uncoupling in brown adipose tissue (BAT) are important modes of heat generation in mammals (Krauchi & Wirz-Justice, 1994; Refinetti & Menaker, 1992a; Nicholls & Locke, 1984; Cannon & Nedergaard, 2004; Refinetti & Menaker, 1992b), which are elevated during wakeful activity that typically occurs at night in mouse, and daytime in humans. Increased peripheral blood flow is the primary mechanism regulating heat loss in mammals, and in humans is essential for lowering core body temperature at night around the transition from wake to sleep (Aschoff & Heise, 1974; Smolander *et al*, 1993; Murphy & Campbell, 1997; Rzechorzek *et al*, 2022). Our observations are consistent with the hypothesis that, *via* NO signalling, RBC metHb rhythms contribute to increased peripheral blood flow and vasodilation during the rest phase, which would allow core body and brain temperature to cool and so facilitate the transition to sleep. Indeed, altered rhythms of body temperature in tau mutant hamsters are consistent with the predicted contribution of cell autonomous metHb rhythms to cooling (Refinetti & Menaker, 1992b). We tested a prediction from this hypothesis, that elevated metHb will accentuate the daily drop in core body temperature (Figure 4). We found that treatment of mice with nitrite, which further increases metHb levels (Figures 1H and 4B), led to a small but significant reduction in body temperature (Figure 4D) in mice during the daytime.

### RBCs and the TTFL-less clock mechanism

Recent observations have increased focus on post-translational events in signalling biological timing information under a putative post-translational oscillator model (Wong & O’Neill, 2018; Crosby & Partch, 2020). The PTO model is compatible with nucleate and anucleate cell types and, accordingly, roles for the clock-related kinases CK1, CK2 and proteasomal degradation have been demonstrated in RBCs (Beale *et al*, 2019; Cho *et al*, 2014). Here, through choosing mutations in proteins that are not expressed in RBCs from humans (Beale *et al*, 2019; Bryk & Wiśniewski, 2017), we further delineated the RBC clock mechanism from PER/CRY-regulated rhythms found in other cell types. Our novel reporter of RBC circadian rhythms, which relies solely on cell lysis and gel electrophoresis without antibodies, will greatly aid the further teasing apart of the post-translational clock mechanisms of RBCs.

## Material and Methods

### Experimental animals and human research participants

All mouse work was licensed under the UK Animals (Scientific Procedures) Act of 1986 with local ethical approval (details given below) and reported in accordance with ARRIVE guidelines. Studies involving human participants were conducted in accordance with the principles of the Declaration of Helsinki, under informed consent, with ethical approval from local research ethics committee.

### Mouse RBC collection for *in vitro* experiments

Mouse work was overseen by the Animal Welfare and Ethical Review Body of the MRC Laboratory of Molecular Biology. Mice were entrained under 12:12 h light:dark cycles from birth prior to being killed at Zeitgeber time 4 (ZT4) by a schedule one method and exsanguinated by cardiac puncture. Blood was immediately transferred to a collection tube (Sarstedt, Nürnbrecht, Germany) containing sodium citrate anti-coagulant, with approximately 8 ml total blood being collected and pooled from 10 isogenic animals (C57BL/6 background, aged 3-4 months) with equal numbers of males and females for each genotype. RBCs were isolated from anticoagulant-treated whole blood by density gradient centrifugation and washed twice in PBS before resuspension in modified Krebs buffer (KHB, pH 7.4, 310 mOsm to match mouse plasma) as described previously (O’Neill & Reddy, 2011). RBC suspensions were aliquoted into single 0.2 mL PCR tube per time point per mouse and incubated at constant 37°C. At each time point the red cell pellet was resuspended by trituration and 50 μl was removed and added to 450 μL 2x LDS sample buffer (Life Technologies, Carlsbad, CA) supplemented with 5 mM DTPA as described previously (Milev *et al*, 2015).

### Human RBC collection for *in vitro* experiments

Studies were conducted with ethical approval from the Local Research Ethics Committee (Cambridge, UK). Participants in the study were screened for health (by history, physical examination, and standard biochemistry and haematology), and did not suffer from sleep disorders or excessive daytime sleepiness. All participants provided written, informed consent after having received a detailed explanation of the study procedures. ∼10 ml of blood was collected from each subject in the morning (8-10AM) using tubes containing sodium citrate anticoagulant (Sarstedt). Red cell pellets were obtained using the same method described above, except that Krebs buffer osmolarity was adjusted to 280 mOsm/L to match conditions normally found in human plasma. 100 µl of re-suspended RBCs were dispensed into 0.2-ml PCR tubes (Thermo), and then placed in a thermal cycler (Bio-Rad Tetrad) at constant 37°C for time course sampling, or else maintained at room temperature for immediate experimentation, as described in the text. At each time point the red cell pellet was resuspended by trituration and 50 μl was removed and added to 450 μL 2x LDS sample buffer (Life Technologies, Carlsbad, CA) supplemented with 5 mM DTPA as described previously (Milev *et al*, 2015).

### Mouse blood collection for *in vivo* sodium nitrite treatment

Mouse work was overseen by the Animal Welfare and Ethical Review Body of the MRC Laboratory of Molecular Biology. C57BL/6J mice (n=12, 6 females, 6 males, aged 9 weeks at the start of the experiment) were singly housed in cages containing environmental enrichment and running wheels for monitoring animal health in two light-tight cabinets. Mice were included in the study if daily locomotor activity pattern was consistent, weight throughout was 20g ± 2g females and 26g ± 2g males, and excluded if mice were inactive for more than 6h during a dark period during recording. The two cabinets were set to opposite 12h:12h light:dark cycles, i.e., animals were oppositely entrained. On day 8, mice were split randomly into treatment and control groups in even sex ratios by mouse ID, splitting littermates evenly between groups. N=4 (2 females, 2 males) mice were given 12.5 mg/kg sodium nitrite (52mg/L, 23713, Sigma) in 10% blackcurrant squash (Robinsons); control mice (n=8, 4 females, 4 males) were given blackcurrant squash. On day 18, all mice were culled at the same external time but different zeitgeber times (ZT), i.e., ZT3 (n=4 control mice) or ZT15 (n=4 control mice, and n=4 mice with nitrite), where ZT0 represents the start of the light phase, by CO2 with exsanguination by cardiac puncture, and whole blood was transferred into tubes containing sodium citrate anti- coagulant. An equal amount of whole blood from each mouse, normalised by Hb concentration as measured by absorbance at the isosbestic point of Hb and metHb (529 nm) using a TECAN plate reader, was lysed by adding an equal volume of 4X LDS 10 mM DPTA to a final concentration of 2X LDS 5 mM DPTA, triturating 4 times, and incubating at room temperature for 15 min. Blood was heated for 10 min at 70°C on a shaking heat block, left to cool then flash frozen. Lysed blood was subsequently diluted to 1X LDS and 50 mM TCEP for gel electrophoresis. Sample size calculation was based on *in vitro* RBC timecourse data (peak mean = 50, trough mean = 30, standard deviation = 10). The power of the experiment was set to 80%. A total of 4 mice per group were considered necessary.

### Human whole blood collection in controlled laboratory conditions

Study procedures were approved by the Institutional Review Board at the University of Colorado Boulder and subjects gave written informed consent. Participants (N=4 [2 male, 2 female], aged 27.3 ± 5.4 years [mean ± SD]) were healthy based on medical and sleep history, physical exam, normal BMI, 12-lead electrocardiogram, blood chemistries, clinical psychiatric interview, polysomnographic sleep disorders screen, and provided written, informed consent. Participants were excluded from study for current or chronic medical or psychiatric conditions, pregnancy, shift work, or dwelling below Denver altitude (1600 m) the year prior to study. Travel across more than one time zone in the three weeks prior to the laboratory study was proscribed. Participants were medication and drug free based on self-report and by urine toxicology and breath alcohol testing upon admission to the laboratory. Participants maintained consistent 9h sleep schedules for at least one-week prior to the laboratory protocol verified by wrist actigraphy and call-ins to a time stamped recorder. Following two baseline laboratory days, participants were awakened at habitual waketime and studied under dim light conditions (< 10 lux in the angle of gaze during wakefulness and 0 lux during scheduled sleep), given an energy balanced diet (standardised breakfast, lunch and dinner meals given at 0800, 1400 and 1800 respectively, and two after dinner snacks at 2000 and 2200h, water available *ad libitum* for someone with a 0600h waketime). Blood samples (10 ml) were collected every two hours from each of the four participants into lithium heparin Vacutainers (BD Biosciences, Franklin Lakes, NJ) and separated by centrifugation. Cell fractions were flash frozen in liquid N2 and stored a -80°C until lysis with 2x LDS sample buffer.

### Gel electrophoresis and immunoblotting

Western blotting and SDS-PAGE were performed, under reducing conditions, as described previously (Milev *et al*, 2015), using antibody against PRX-SO2/3 (Abcam, ab16830). For bloody blotting, membranes were quickly washed with deionised water and then immediately incubated with ECL reagent (Millipore). Coomassie stained gels were imaged using an Odyssey scanner (Licor). Chemiluminescence was detected by x-ray film, except for the data in Figure 2D-G which were collected on a ChemiDoc XRS+ (Bio-rad).

### Protein digestion and mass spectrometric analysis

Peptides were prepared from excised bands from Coomassie stained gel as follows. Gels slices were first destained in 50% acetonitrile/50% 100mM NH4HCO3. After washing in 2 times 100μl dH2O, gel slices were dehydrated by the additional of 100μl of acetonitrile, and the solvent completely evaporated by lyophilising. Proteins within the bands were first reduced by the addition of 50μl 10mM dithiothreitol in 100mM NH4HCO3 and incubated at 37°C for 1 hour. The supernatant was then removed and replaced by 50μl of 55mM iodoacetamide in 100mM NH4HCO3 and incubated at room temperature in the dark for 45 minutes. After removal of the supernatant the bands were washed in 100mM NH4HCO3 followed by 100% acetonitrile prior to dehydration as described above.

Trypsin lysis was carried out by first rehydrating the bands in 300ng trypsin (Sequencing Grade Modified Trypsin (Promega Product no. V5111) in 60μl 100mM NH4HCO3 and incubated overnight at 37°C. The supernatant was then acidified to 0.1% formic acid. Peptide analysis was performed by matrix assisted laser desorption ionisation (MALDI) mass spectrometry (Waters Micromass MaldiMX Micro) using a-cyano-4-hydroxycinnamic acid matrix (10 mg ml−1 in 50% aqueous acetonitrile/0.1% trifluroacetic acid).

### Hb biochemistry

All reagents were purchased from Sigma-Aldrich (St Louis, MO) unless otherwise stated. Direct staining of sodium dodecyl sulfate (SDS)-containing polyacrylamide gels with o-dianisidine, to detect protein-bound to haem was performed as described previously (Maitra *et al*, 2011). All RBC incubations in Figure 2 were performed in Krebs buffer. All stock solutions were prepared at 2 M in deionised water except N-ethylmaleimide, which was prepared in ethanol. Native purifications were performed using Ni-NTA agarose according to (Ringrose *et al*, 2008), denaturing purifications in 8M urea were performed according to manufacturer’s instructions.

### Bioluminescence recording

Primary fibroblasts carrying a *Cry1:luciferase* reporter (Maywood *et al*, 2013) were isolated from the lung tissue of adult males (C57BL/6) and cultured as described previously (O’Neill & Hastings, 2008), and verified mycoplasma free by Lonza MycoAlert mycoplasma detection kit (LT07, Lonza). Fibroblasts were synchronized by temperature cycles of 12 h 32°C followed by 12 h 37°C for 5 days, then changed to ‘‘Air Medium’’ (Bicarbonate-free DMEM (D5030, Sigma) dissolved in dH2O, modified to a final concentration of 5mg/mL glucose, 0.35 mg/mL sodium bicarbonate, 20 mM MOPS, 2 mg/mL pen/strep, 2% B27, 1% Glutamax, 1mM luciferin, and adjusted to pH 7.6 and 350 mOsm; adapted from (O’Neill & Hastings, 2008)), and dishes sealed. Bioluminescence recordings were performed using a Lumicycle (Actimetrics, Wilmette, IL, USA) under constant conditions as described previously (Causton *et al*, 2015). Lumicycle data were detrended to remove baseline changes and then fit with a damped sine wave to determine circadian period as in (Causton *et al*, 2015).

### Pulse co-oximetry

Pulse co-oximetry was performed using a Masimo Radical 7 (Masimo, Irvine, CA, USA) according to manufacturer’s instructions using a finger probe to measure blood parameters in the periphery. Co-oximetry records pulse rate (PR), perfusion index (PI, the ratio of the pulsatile blood flow to the nonpulsatile static blood flow in a peripheral tissue), and the proportion of total Hb (SpHb), Oxygenated Hb (SpO2) and MetHb (SpMet) in the blood. Four freely-behaving, healthy, age-matched male volunteers (25-35 years old) were monitored over a 3-day period during a normal working week.

Data was collected in Cambridge, UK between September and November, 2012. Data collection was conducted in accordance with the principles of the Declaration of Helsinki, with approval/favourable opinion from the Local Research Ethics Committee (University of Cambridge, UK). Participants in the study were screened for relevant self-reported health issues, including sleep disorders or excessive daytime sleepiness.

### Mouse body temperature measurement

Animal experiments were approved by the animal welfare committee at the University of Manchester. C57BL/6J mice were purchased from Charles River (UK). N=8 mice were housed under a 12:12 light:dark cycle at ∼400:0 lux for the duration of the experiment. Ambient temperature was maintained at 22± 2 °C and humidity at 52 ± 7%, with food and water, or 10% blackcurrant squash (Morrisons) ± drug, available *ad libitum*. Mice were group housed until the start of experimental recordings when they were individually housed, and kept in light-tight cabinets. Male mice aged 9 weeks at the beginning of the experiment were used.

Mice were implanted with TA-F10 radiotelemetry devices (Data Sciences International, USA) to record body temperature and locomotor activity. Radiotelemetry devices (sterilised in 2.5% Gluteraldehyde (G6257, Sigma) and rinsed with sterile saline) were implanted in the abdominal cavity of isofluorane anaesthetised mice (2-5% in oxygen, total duration ∼15 minutes), followed by a recovery period of 10 days. Body temperature and locomotor activity were recorded every 5 minutes throughout the experimental recording period. Locomotor activity monitoring was used to monitor mouse health during the experiment. Mice were included in the study if healing after implantation was complete, and no wound opening was observed. Mice were excluded if surgical wounds opened at any point during the experiment, or locomotor activity was absent for more than 6h during recording.

After a 2-day habituation period to the individual cages, baseline recording was collected for 2 full days. At ZT23, after the 2 full days of baseline recording and prior to the inactive period of mice, mice were given sodium nitrite (375mg/L, 23713, Sigma) diluted in 10% squash for palatability for 2 further full days with body temperature and locomotor activity recorded. To generate average day profiles during baseline or sodium nitrate phases of the experiment, 15-minute bins of body temperature were averaged across the two days of recording for each mouse. The average daily profile of all mice was plotted against time of day. To quantify effect of nitrite on body temperature, controlling for locomotor activity and time of day, each 15-minute bin of body temperature was categorised as belonging to ‘active’ or ‘inactive’ locomotor periods based on whether locomotor activity during the bin was higher or lower, respectively, than the individual’s median activity across the whole 2-day experimental recording period. The median threshold was chosen to keep the number of ‘active’ and ‘inactive’ bins similar, though a more stringent threshold of inactivity (total inactivity across 3 consecutive bins) was also assessed, and similar results observed. Body temperature in bins belonging to ‘active’ and ‘inactive’ periods were separated by time of day (light, day vs dark, night) and averaged for each mouse across the two days of recording. Thus, for each mouse, four separate average body temperatures were generated, belonging to inactive and active periods during the light phase, and inactive and active periods during the dark phase. Average body temperatures for the ‘inactive’ and ‘active’ periods in light and dark phases were compared across baseline and sodium nitrite phases of the experiment by repeated measures three-way ANOVA.

### Statistical Analysis

All graphs and analyses were performed in Prism 8 (Graphpad Software Inc, La Jolla, CA) and R (version 3.6.3) in R Studio (version 1.2.5033, RStudio Inc). Least squares non-linear regression curve fitting in GraphPad Prism (v7.03, GraphPad; La Jolla, CA) was used to determine whether circadian rhythms were present in quantified data by comparing straight line fit and damped cosine fit with the extra sum-of-squares F test as described previously (Beale *et al*, 2019). Pulse co-oximetry data was analysed for significant rhythmicity by autocorrelation in R and subsequent damped cosine curve fitting as above. Mean ± SEM is reported throughout. Analyses are reported in figure legends.

## Data Availability

This study includes no data deposited in external repositories.

## Acknowledgments

We are grateful to the LMB Biomedical Services Group for animal care, the donors and Noel Wardell for assistance with the blood samples, Kevin Feeney, Guillaume Rey, and Nina Rzechorzek for useful discussion. We also thank Mark Skehel for assistance with mass spectrometry.

## Author contributions

JON and MHH designed experiments; JEC, ESM, ADB and EAH performed animal work; PC performed fibroblast assays, RSE, JON performed RBC time courses and biochemistry; ADB performed acute whole blood biochemistry; ABR performed phlebotomy; KSL performed mass spectrometry; UV, JON and ABR collected and analysed co-oximetry data; ADB performed data analysis; FHL, ABR, and DAB contributed resources and discussion; KPW collected human blood samples in controlled laboratory conditions; MHH and JON supervised the work; ADB & JON wrote the paper.

## Expanded View Figure Legends

**Figure EV1.**
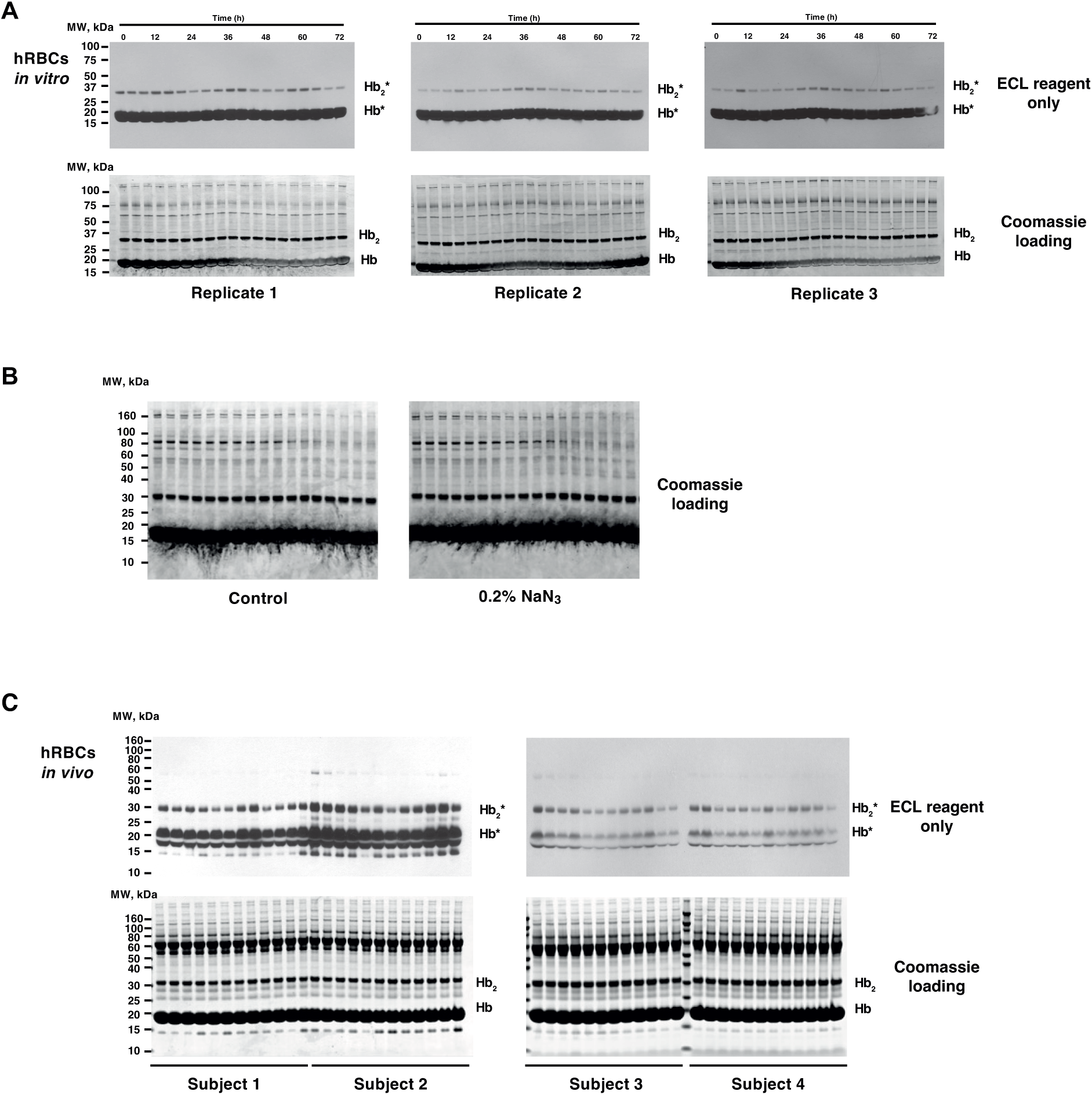
relating to Figure 1 and Figure 3 - Daily variation of haem-mediated peroxidase activity in human RBC time course extracts on nitrocellulose membranes following SDS-PAGE under denaturing conditions. **A** Uncropped blots and gels from in vitro human RBC time course extracts shown in Figure 1. Top panels show the chemiluminescence signal arising when nitrocellulose membranes were immediately incubated with ECL reagent, following SDS-PAGE and transfer onto nitrocellulose membranes. Please note that chemiluminescence is observed at molecular weights corresponding to Hb and Hb2 without any antibody incubation or external source of peroxidase activity. For practical reasons, the Hb2* band was used for quantification presented in Figure 1, since Hb* quickly saturated the x-ray film used to detect chemiluminescence. Bottom panels show associated coomassie-stained gels (loading control). **B** Uncropped coomassie-stained gels from in vitro human RBC time course extracts shown in Figure 1B. In Figure 1B, before incubation with ECL reagent, membranes were incubated for 30 minutes in PBS ± 0.2% sodium azide. **C** Uncropped blots and gels from human blood time course sampled in vivo shown in Figure 3. Top panels show the chemiluminescence signal arising when nitrocellulose membranes were immediately incubated with ECL reagent, following SDS-PAGE and transfer onto nitrocellulose membranes. Please note that chemiluminescence is observed at molecular weights corresponding to Hb and Hb2 without any antibody incubation or external source of peroxidase activity. For practical reasons, the Hb2* band was used for quantification presented in Figure 3, since Hb* quickly saturated the x-ray film used to detect chemiluminescence. Bottom panels show associated coomassie-stained gels (loading control).

**Figure EV2.**
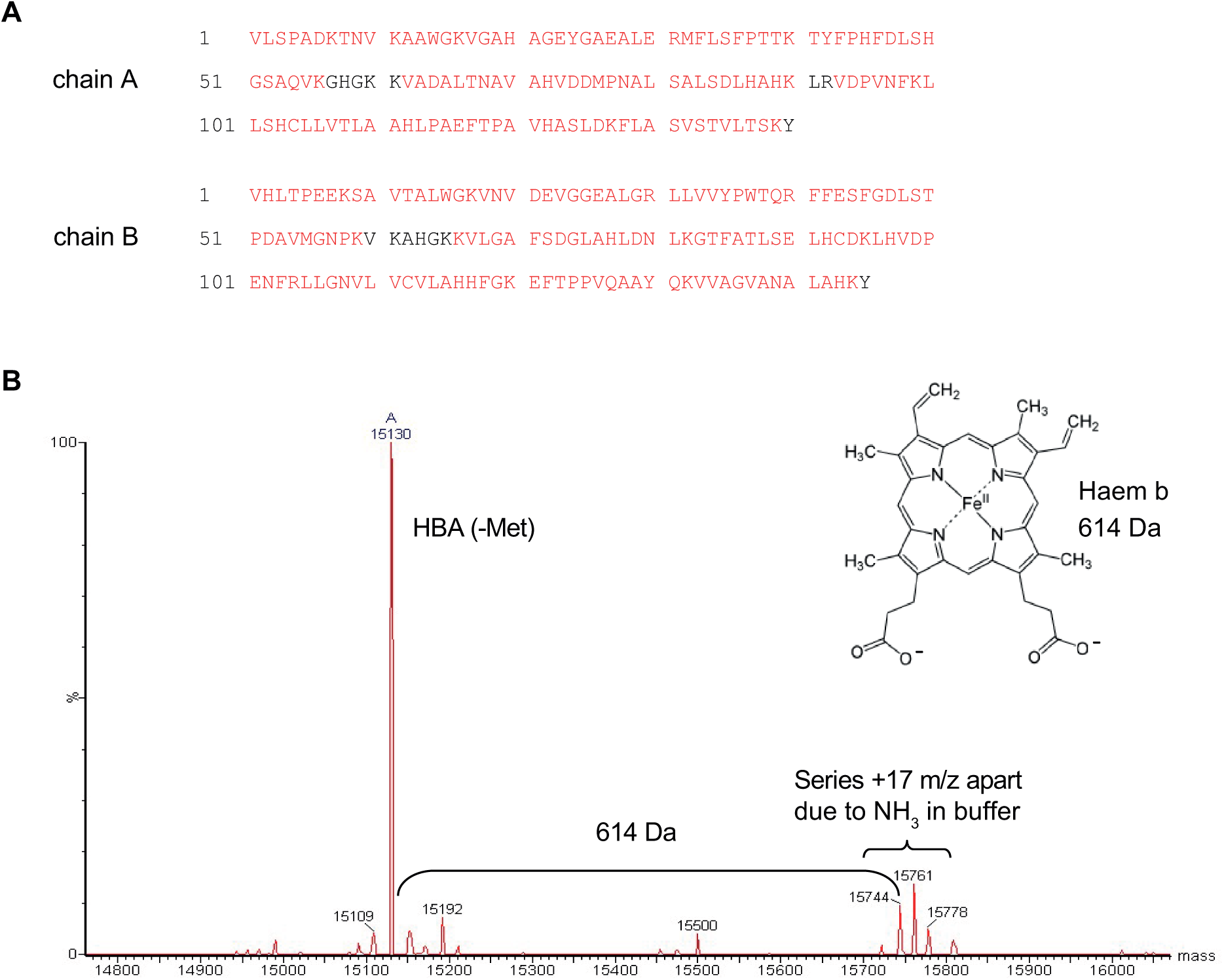
relating to Figure 1 - Mass spectrometry of Ni-NTA purified peroxidase band. **A** Mass spectrometry of tryptic digests of the putative haem-Hb species purified by Ni-NTA affinity chromatography under denaturing conditions (Figure 1C) revealed Hb chain A and B (HBA/HBB) as the major species with very high primary sequence coverage (indicated in red). **B** The haem-Hb species purified by Ni-NTA affinity chromatography under denaturing conditions was excised and the undigested polypeptide analysed by mass spectrometry. Raw mass spectrometry data showing signal at the expected molecular weight of HBA protein are shown, with additional peaks at +614 Da - the mass of haem b (insert).

**Figure EV3.**
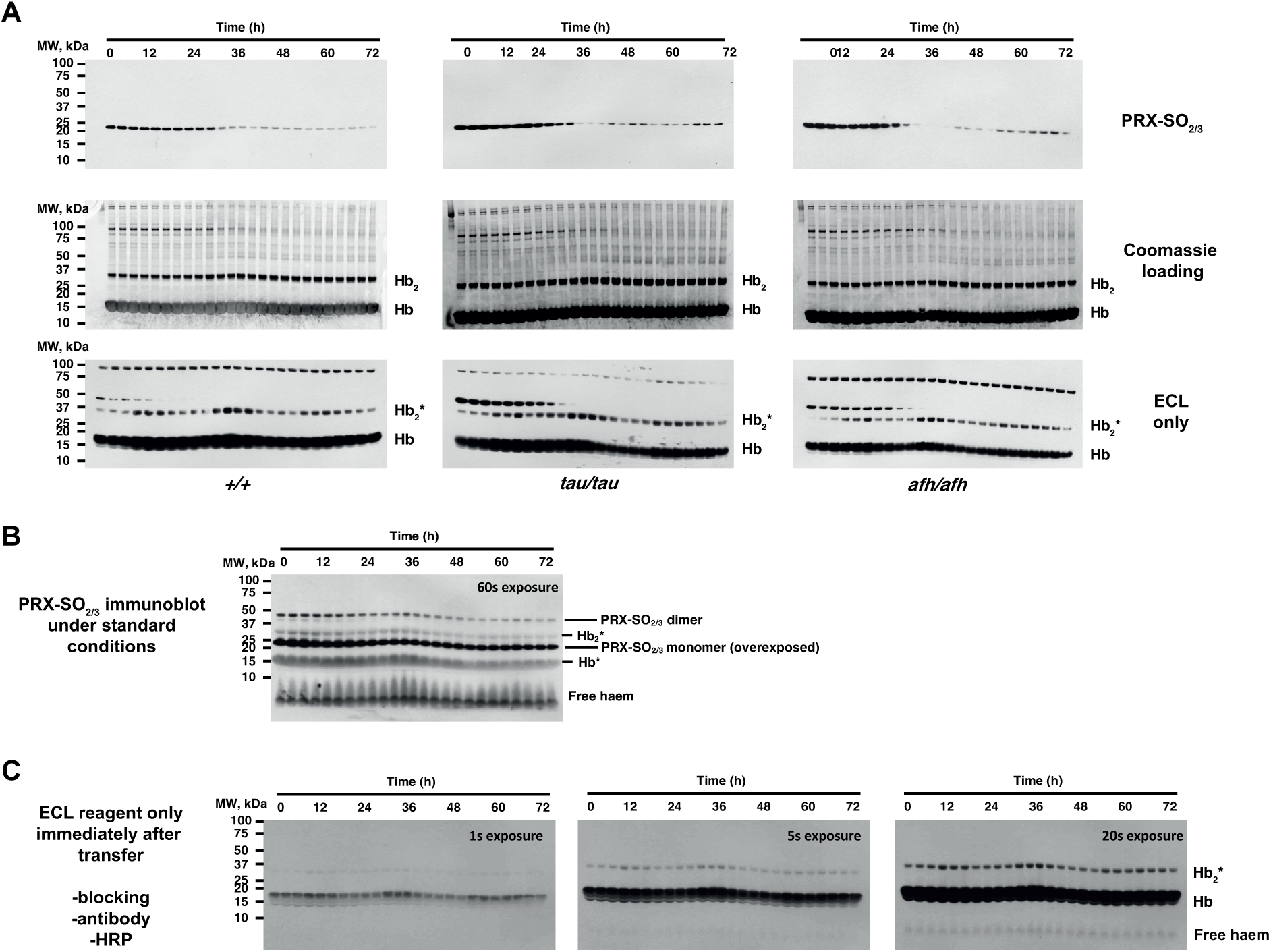
relating to Figure 2 - Circadian variation of PRX-SO2/3 and haem-mediated peroxidase activity in mouse RBC time course extracts on nitrocellulose membranes after denaturing SDS-PAGE transfer. **A** Top panels show uncropped representative PRX-SO2/3 immunoblots of RBC time course extracts from each genotype shown in Figure 2. Middle panels show associated coomassie-stained gels (loading control) where Hb is the major protein. The previously reported occurrence of cross-linked Hb dimers (Hb2) is readily observable at ∼32 kDa, and the identity of the band marked as Hb2 was confirmed by mass spectrometry (not shown). Bottom 3 panels show the chemiluminescence signal arising when replicate nitrocellulose membranes were immediately incubated with ECL reagent, following transfer from SDS-PAGE. Please note that chemiluminescence is observed at molecular weights corresponding to Hb and Hb2 without any antibody incubation or external source of peroxidase activity. For practical reasons, the Hb2* band was used for quantification presented in Figure 2, since Hb* quickly saturated the x-ray film used to detect chemiluminescence (as shown in C, below). **B** The intrinsic chemiluminescence from bands at molecular weights corresponding to Hb and Hb2, as well as free haem, was first observed faintly as non-specific bands in over-exposed PRX-SO2/3 immunoblots. **C** Further investigation revealed that this peroxidase activity was not attributable to non-specific antibody binding, since is readily observed in RBC extracts upon addition of ECL reagents to nitrocellulose membranes, immediately after transfer from SDS-PAGE. Please note the very high activity of Hb* compared with Hb2*, which is consistent with the relative levels of Hb and Hb2 detected by coomassie in (A). Also, note the signal due to free haem that is apparent upon longer exposures (right). Interestingly, compared with human RBC time courses (Henslee et al., 2017; O’Neill and Reddy, 2011), we observed that murine PRX-SO2/3 immunoreactivity was extremely high during the first 24 hours of each 72-hour time course (Figure S3A). We attribute this to the different conditions under which blood was collected: blood was collected from mice culled by CO2 asphyxiation during their habitual rest phase by cardiac puncture and exposed immediately to atmospheric oxygen levels, whereas human blood was collected from subjects during their habitual active phase through venous collection into a vacuum-sealed collection vial. Thus, the initial high PRX-SO2/3 signal in mice may be related to CO2-acidification of the blood during culling, which affects PRX-SO2/3 but does not affect Hb oxidation status.

**Figure EV4.**
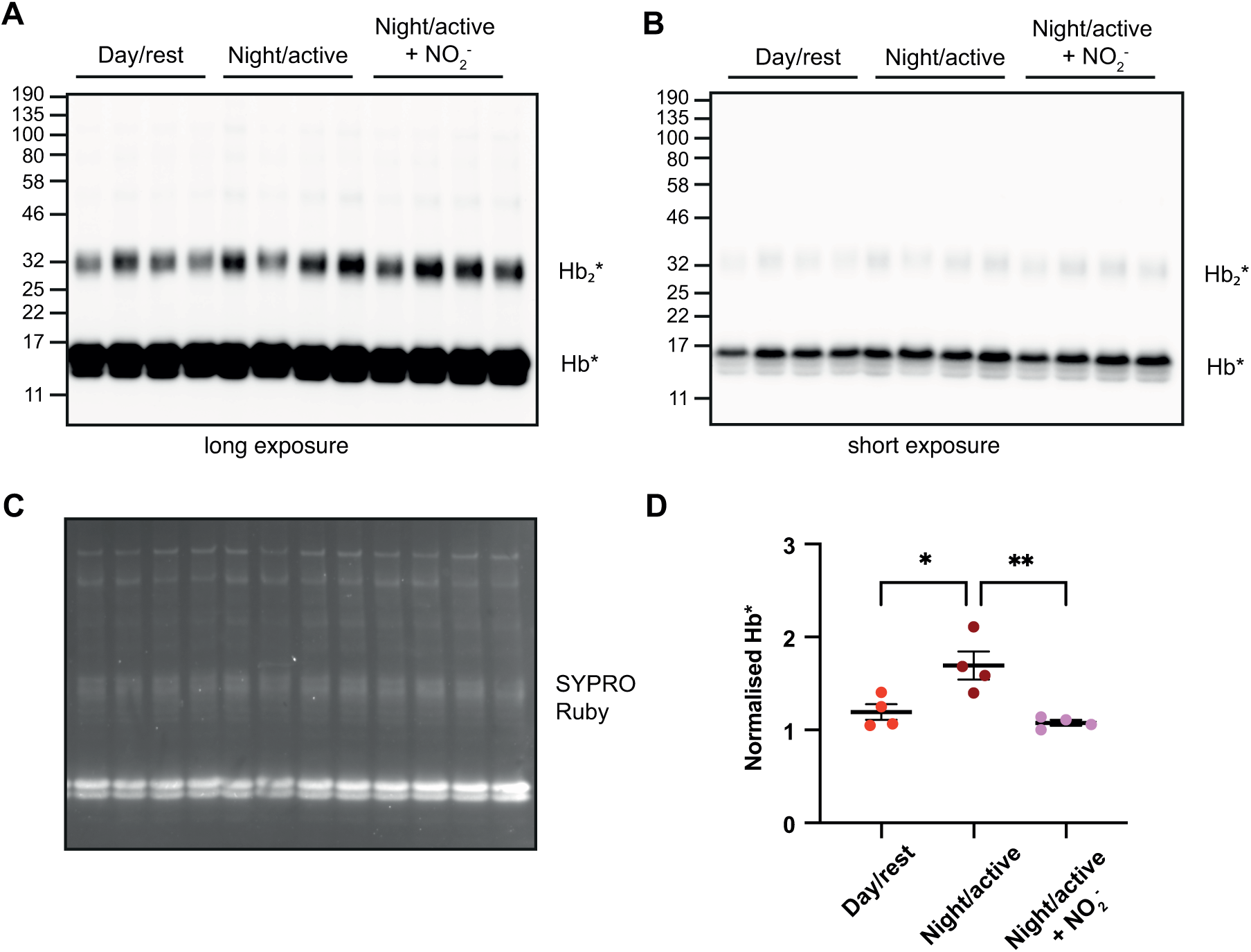
relating to Figure 4 - Uncropped nitrocellulose membranes and gels as presented in Figure 4. **A** Long exposure, for quantification of the Hb2* band as presented in Figure 4. **B** Short exposure, showing Hb* activity. **C** SYPRO Ruby uncropped gel. The lower band of SYPRO Ruby (corresponding to Hb monomer) is presented in Figure 4, but the intensity of the whole lane was used for quantification of loading. **D** Quantification of Hb* band of short exposure, using same methods as Figure 4. The same result is observed when quantifying Hb* as Hb2*.

