## Supplemental Figure 1 for "Mechanisms and physiological function of daily haemoglobin oxidation rhythms in red blood cells"

### Slide 1
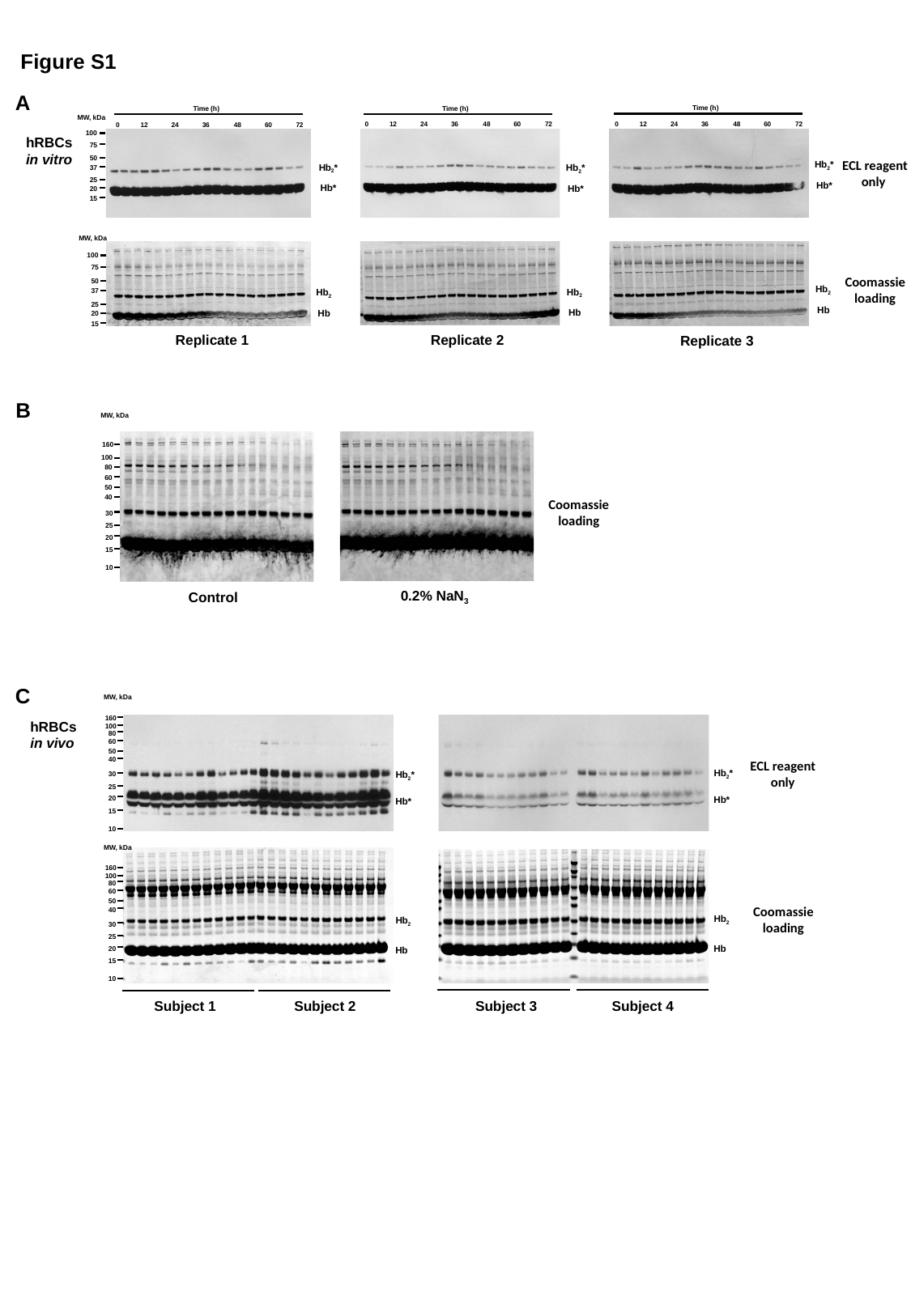

Figure S1
A
Time (h)
0
12
24
36
48
60
72
Time (h)
0
12
24
36
48
60
72
Time (h)
0
12
24
36
48
60
72
MW, kDa
100
hRBCs
in vitro
75
50
ECL reagent
only
Hb_2_*
Hb_2_*
Hb_2_*
37
25
Hb*
Hb*
Hb*
20
15
MW, kDa
100
75
50
37
25
20
15
Coomassie
loading
Hb_2_
Hb_2_
Hb_2_
Hb
Hb
Hb
Replicate 1
Replicate 2
Replicate 3
B
MW, kDa
160
100
80
60
50
40
30
25
20
15
10
Coomassie
loading
0.2% NaN_3_
Control
C
MW, kDa
160
100
80
60
50
40
30
25
20
15
10
hRBCs
in vivo
ECL reagent
only
Hb_2_*
Hb_2_*
Hb*
Hb*
MW, kDa
160
100
80
60
50
40
30
25
20
15
10
Coomassie
loading
Hb_2_
Hb_2_
Hb
Hb
Subject 3
Subject 4
Subject 1
Subject 2
